# Compositional Analysis of Mouse Sperm Subpopulations During Capacitation In Vitro

**DOI:** 10.64898/2025.12.23.696226

**Authors:** Benjamin M. Brisard, Aishwarya P. Halder, Aidan Charles, Amy L. Ward, Maryam Asadi, David Hart, Debajit Bhowmick, Paul Vos, Cameron A. Schmidt

## Abstract

Mammalian sperm must undergo capacitation to become competent for fertilization, yet this process is marked by substantial phenotypic heterogeneity among sperm cells. How such variability emerges and how it relates to fertilizing potential remain unresolved, in part because sperm subpopulations are typically analyzed independently despite being intrinsically interdependent. Here, we combine large-scale single-cell spectral flow cytometry with compositional statistical modeling to quantify how sperm population structure responds to controlled capacitation signals in vitro. Using cauda epididymal mouse sperm, we implemented a two-dimensional assay that systematically varies extracellular bicarbonate and free calcium—key regulators of capacitation—and classified millions of individual cells into four irreversible physiological states defined by cell viability and acrosome reaction status. We show that bicarbonate and calcium interact nonlinearly to redistribute sperm across these subpopulations, revealing structured responses that would be otherwise obscured in measurements lacking single-cell resolution. Elevated intracellular calcium was associated with increased cell death, while the highest proportions of live, acrosome-reacted sperm occurred under relatively low extracellular calcium conditions. To enable subpopulation-level analysis, we applied a hierarchical Dirichlet–multinomial regression model that accounts for multinomial sampling noise and between-male variability, yielding posterior probability surfaces that describe how sperm subpopulations reallocate across functional states as microenvironmental signaling conditions change. Together, these results demonstrate that capacitation is a stochastic, cell-population level process shaped by structured phenotypic heterogeneity. This framework provides a quantitative foundation for linking sperm subpopulation composition to measures of fertility competence and for improving existing interpretation of flow cytometry–based assessments of male fertility.

## Introduction

Mammalian sperm must undergo a post-ejaculatory maturation process, known as capacitation, before they can fertilize an egg. Within the female reproductive tract, sperm encounter biochemical effectors such as bicarbonate or progesterone in the luminal fluid microenvironment, and exposure to these agents *in vitro* reproducibly induce capacitation^1,2^. Capacitation is a multifaceted maturation process, and the complete understanding of its mechanistic constituents remains a matter of ongoing investigation^1^. However, several decades of empirical work implicate a subset of highly conserved biochemical processes including potassium-dependent plasma membrane hyper-polarization^3^, increased intracellular pH^4^, 3’,5’-cyclic adenosine monophosphate (cAMP)-dependent PKA activation^5^, protein tyrosine phosphorylation^6,7^, and a sustained rise in intracellular calcium^8^. This conserved system of regulatory contingencies explains multiple aspects of sperm behavior including motility patterns such as the progressive to hyperactive transition^9^, acrosomal exocytosis (a.k.a. acrosome reaction)^10^, and other putative sperm guidance mechanisms such as chemotaxis and chemokinesis^11^. Other conserved sperm behaviors do not require direct biochemical control, but rather manifest from intrinsic cellular mechanics under the constraint of microscale physical forces, such as collective motion, rheotaxis, and thigmotaxis^12–14^.

Despite ongoing attempts to develop a deterministic molecular control model of sperm capacitation, these changes are often not uniformly or universally achieved within an ejaculate. Significant heterogeneity has been reported at nearly every level of the proposed cell signaling control pathway^15^, including calcium permeability^16,17^, plasma membrane hyperpolarization^18^, kinetics of intracellular ion transients ^19^, protein tyrosine phosphorylation^20^, extent and rate of the acrosome reaction^21^, and metabolic state^2^. This conserved variation highlights that the ejaculate is not a homogenous population of identical cells, but rather a collection of stochastically varying individuals with distinct phenotypes that manifest uniquely depending on microenvironmental conditions. Furthermore, repeated measures sampling of sperm functional parameters from the same males over time also reveals substantial heterogeneity, indicating that sperm phenotypic variation is an extremely complex, multiscale stochastic phenomenon, yet how this complexity relates to fertility competence remains largely unknown^22,23^.

The persistence of sperm heterogeneity raises a central question: why should sperm, whose ultimate physiological function is to fertilize an egg, display such variable responses to common signaling inputs that should ostensibly improve the likelihood that they will be successful? One possibility is that heterogeneity reflects an unavoidable and potentially maladaptive “noise” as a simple consequence of deleterious mutations during meiosis^24,25^ or incomplete maturation during residence in the epididymis^26^. In this scenario, fitness would most likely decrease with increasing variation. However, heterospermic experiments (such as those performed in birds) indicate that the opposite is true, heterogeneity generally increases fitness^27^. In contrast to maladaptive noise, phenotypic heterogeneity among sperm may confer enhanced fitness as an *exploratory mechanism*. Under this model, each sperm may be similar to others, but unique in their exact combination of phenotypic characteristics. Importantly, sperm are transcriptionally and translationally repressed, and as such, may access only a limited repertoire of adaptive behaviors in response to environmental stressors as a function of their genotype^28^.

When large numbers of sperm search for an egg, each with slight variation, the phenotype distribution may enable the cell population as a whole to hedge against environmental uncertainty better than any individual sperm can on its own. In other words, large sperm numbers may have evolved as a way for sperm to off load adaptability under the constraints imposed by transcriptional/translational repression. In this view, heterogeneity is not strictly maladaptive noise, but an essential physiological feature that shapes fertility competence when the conditions of the fertilizing microenvironment are not known to the male a priori. Through statistical variation, an ejaculate may explore a multiplicity of phenotypes in parallel, hedging on the probability that at least some sperm will be competent to achieve fertilization under variable conditions or unknown time to egg availability^11,27^.

Understanding this balance between sperm individuality and collective adaptability may reveal new methods for predicting male fertility from semen parameters, thereby improving accuracy of fertility diagnostics or enhancing IVF efficiency. Under these assumptions, fertility depends on the statistical dynamics of phenotype distributions, necessitating analysis methods that treat sperm subpopulations as dependent wherein a change in the proportion of one subpopulation must reflect a change in another. This approach may augment current ensemble averaging methods^29^, conferring a significant advantage because it does not require any a priori assumptions about what constitutes the ‘best’ sperm^30^—a common but logically dubious approach that ignores the context dependence of fitness^31^.

Developing and testing a generalizable model of sperm exploratory dynamics will require new experimental methods and statistical tools that correlate fertility with the dynamics of sperm subpopulation compositional change. Flow cytometry has proven extremely useful for single-cell assessment of sperm phenotype distributions and played a key role in the initial discovery of sperm subpopulations^15,17,32–34^. Here, we develop a novel flow cytometry approach to measure and model sperm subpopulation compositional responses across millions of individual sperm—isolated from the cauda epididymis of mice. We use intracellular calcium, acrosome reaction status, and cell viability as surrogate measures of phenotypic changes related to in vitro capacitation. We employ a novel method of systematic cell signaling control through 2D scaling of a capacitive signaling input (bicarbonate) and an essential second messenger (extracellular calcium), as well as their interaction (bicarbonate x calcium). We then apply phenotypic classification and a hierarchical regression modeling methods originally developed for compositional analysis of taxonomic distributions in microbial ecology to study how sperm subpopulation proportions respond to microenvironments with a range of signaling impulses.

## Materials and Methods

Chemicals and reagents were sourced from Sigma-Aldrich (St. Louis, MO). Specific culture media were employed to facilitate various experimental conditions. Human tubal fluid minimal media (HTF-min) was used as a base medium with (in m*M*) sodium chloride (91.2), potassium phosphate monobasic (0.37), magnesium sulfate heptahydrate (0.20), potassium chloride (4.69), sodium L-pyruvate (0.33), sodium L-lactate (21), and D-glucose (2.78), Bovine serum albumin (BSA) (4mg/mL), Sodium 4-(2-hydroxyethyl)-1-piperazineethanesulfonic acid (10.4) (HEPES), and HEPES free acid (14.6). pH was maintained at 7.4 at 37°C and stability was verified using a pH microelectrode. Additional supplements such as ethylene glycol bis(2-aminoethyl ether)-N,N,N’,N’-tetra-acetic acid (EGTA) (1), sodium bicarbonate (0-10), and calcium chloride (0-1.8), were incorporated for specific purposes. Media were vacuum filtered through a PTFE membrane with 0.22 um pore size and stored under sterile conditions.

### Animals

Adult male outbred CD-1 retired breeder mice were selected for this study due to having known fertility status. These mice were obtained from Charles River (Wilmington, MA, USA) and received care in accordance with the guidelines set by the National Research Council Guide for the Care and Use of Laboratory Animals. The Institutional Animal Care and Use Committee of East Carolina University approved all experimental procedures. Mice were maintained on a 12-hour light/dark cycle and had access to water and food ad libitum.

### Sperm Isolation

Testes with intact epididymides were dissected into pre-warmed phosphate-buffered saline (PBS, 37°C). Cauda epididymides were transferred to non-capacitating HTF medium lacking bovine serum albumin, sodium bicarbonate, and calcium chloride, and gently dissected to allow sperm to swim out (∼15 minutes at 37°C in a CO_2_ incubator). Residual tissue was removed by centrifugation at 100 x g, and sperm were collected by centrifugation at 800 x g and washed. Sperm counts were obtained using a hemocytometer.

### Validation of Subcellular Dye Localization

Dye localization was confirmed by Laser confocal microscopy (LSM800) with a 40x Plan-Apochromat 1.4 oil DIC 1.4 NA objective. Images were captured in Zen Lite (Zeiss, Jena Germany), background-corrected, converted to 8-bit with an applied 16-color LUT for each composite channel in Fiji ImageJ^35^.

### Validation of the EGTA-Calcium Clamp

Media were prepared with ultrapure water to minimize baseline free calcium. 1 mM EGTA (ethylene glycol-bis(β-aminoethyl ether)-N,N,N′,N′-tetraacetic acid) was used to clamp the ‘free’ Calcium ion concentration in assay preparations. Titrations were measured using reference and calcium ion selective electrodes (Kwik-Tip series; World Precision Instruments (WPI), Sarasota FL). To capture the dynamic range of the media, two standard curves were generated using either 0.1 M CaCl_2_ filling solution or 0.001 M. Once determined, the calcium buffered conditions were included in assay incubations to ‘clamp’ the free calcium within a desired range in conjunction with sodium bicarbonate pseudo-titrations. Linear equations obtained from the standard curves were used to interpolate free calcium concentrations using various combinations of EGTA (1mM) and CaCl_2_. Because free calcium concentration is pH dependent a mixture of HEPES free acid and base were used to buffer pH, which was monitored using a pH microelectrode (WPI).

### Microtiter Fluorometry

Fluorescence measurements were performed using an ID3 Max plate reader (Molecular Devices, San Jose, CA) equipped with Spectramax acquisition software. A two-dimensional 96-well format was designed to maximize the number of experimental conditions evaluated over time. Each condition was monitored at 5-minute intervals, striking a balance between sufficient temporal resolution and minimizing light exposure. Three fluorescent dyes were employed to monitor key physiological parameters: SNARF-1 Acetoxy Methyl Ester (AM; 10 µM) for intracellular pH, Fura-2 AM (10 µM) for intracellular calcium, and Ethidium Homodimer-1 for nuclear membrane permeability as a measure of cell viability. This multiplexed approach enabled time-resolved tracking of sperm responses across a matrix of extracellular calcium and bicarbonate concentrations. All raw fluorescence data were analyzed using GraphPad Prism version 10.4.1.

### Flow Cytometry

For high-resolution single-cell fluorescence measurements, data were acquired on a Cytek Aurora equipped with 405, 488, 561, and 637 nm lasers with SpectroFlo V2.2 acquisition software (Cytek, Fremont, CA, USA). Side scatter (SSC), side scatter–blue (SSC-B), and forward scatter (FSC) gains were optimized to distinguish sperm from debris. Singlet sperm were gated using: Forward scatter Area (FSC-A) vs. Forward scatter height (FSC-H) and Side scatter from the near-UV laser (SSC-B) vs. forward scatter (FSC) to exclude doublets, fragments, and media particles (Initial gating and data export was performed with FlowLogic V8.7). Positive controls were treated for 15 minutes prior to acquisition. The detergent digitonin was used to induce plasma membrane permeability (death) and the calcium ionophore A23187 (10 µM) to induce calcium permeability and stimulate maximal acrosomal exocytosis. The multiplexed dye panel included Indo-1, conjugated peanut agglutinin lectin (PNA)-FITC (1 µg/mL), and TO-PRO-3 (0.1 µM) for simultaneous Ca^**2**+^, acrosome, and viability measurement respectively. Notably, PNA-FITC fluorescence will increase in live mouse sperm as the acrosome reaction progresses due to the presence of externalized inner-leaflet glycoproteins to the outer surface of the plasma membrane^36^. This is sometimes confused with the opposite interpretation due to FITC signal being lost following the acrosome reaction in sperm extracted with polar organic solvent. For solvent extracted sperm, the lectin protein binds to the outer leaflet glycoproteins of the acrosomal vesicle made available by solvent permeation of the plasma membrane; thus the signal decreases following acrosomal exocytosis. Raw data exported from FlowLogic were analyzed using a bespoke program written in Python (V3.11)^37–39^. Source code for the program is publicly available at (https://github.com/CAS-ReproLab/P009_Flow-Cytometry-Tools).

### Classification into Subpopulations

Positive and negative control conditions were included at time of data collection for every assay. For cell viability measured via TO-PRO-3, the positive control condition used digitonin to permeabilize the sperm. For acrosome reaction measured via FITC conjugated peanut agglutinin lectin, the calcium ionophore A23187 was used as a positive control to stimulate the acrosome reaction. A kernel density estimation (KDE) -naive Bayes classifier was used to classify the cells as live/dead or acrosome reacted/unreacted using experimental fluorescence intensity values and control data. Briefly, KDE’s were fit to the fluorescence intensity distributions using Gaussian KDE methods from the Python scikit-learn library^38^. The KDE’s estimate the class conditional probability densities *p(x* | *C*_*k*_), where *x* is the observed fluorescence intensity and *C*_*k*_ is the class. Scott’s rule was used to calculate the bandwidth^40^. The posterior probability was then calculated for experimental data in each filtered treatment condition as follows.

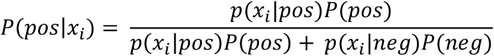

Where lowercase *p* denotes probability density (fluorescence intensity) and uppercase *P* denotes the prior probability for positive (pos) or negative (neg) classes respectively. Prior probabilities were chosen based on review of previous literature and descriptive analysis of the data. The decision rule (*τ)* assigns positive if the posterior probability > *τ*, otherwise assigns negative. The decision rule was chosen for each biological replicate (mouse) by plotting the unfiltered posteriors and manually identifying the mid-point trough in the distribution which was bimodal in all cases.

### Statistical Modeling and Analysis

#### Model summary and motivation

All analyses were performed in Python (v 3.11) using the PyMC probabilistic programming framework (v4.0)^41^. Posterior inference diagnostics and visualization were performed using ArviZ (v0.22.0)^42^. The flow cytometry experiments in this report generate cell counts for mutually exclusive subpopulations of sperm. The data are compositional and inherently constrained, meaning that an increase in one subpopulation’s share of the total population must be compensated by a decrease in another. Modeling approaches commonly used in flow cytometry studies treat cell subpopulations independently, violating this important constraint. To avoid this issue and take full advantage of the power of single-cell measurements, we employed a Dirichlet-Multinomial hierarchical regression model adapted from recent advances in compositional analysis of cell subpopulations in microbial ecology^43–45^. This approach accounts for 1) multinomial sampling noise, 2) biological heterogeneity among replicate animals (overdispersion), and 3) treatment dependent effects on subpopulation composition.

#### Model Overview

Let *y*_*i,k*_ denote the number of cells assigned to subpopulation *k* in 2D assay well *i* (i.e., treatment condition), with total sampled count from the assay well *N*_*i*_ = ∑_*k*_ *y*_*i,k*_. The underlying true subpopulation proportions for that assay well are denoted by the vector π_*i*_ =(π_*i*,1_, …, π_*i,K*_), where *K* = 4 (Live acrosome reacted- L_R_; Live acrosome unreacted- L_UR_; Dead acrosome reacted- D_r_; and Dead acrosome unreacted- D_UR_). We modeled the observed counts as

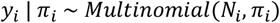

With a Dirichlet prior on the composition:

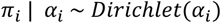

where *α*_*i*_ =(*α*_*i* 1_, …, *α*_*i,K*_ ) is a vector of Dirichlet concentration parameters. Large values of *α*_*i,K*_ correspond to more stable (less variable) subpopulation proportions, whereas small values correspond to greater well-to-well variability around the expected proportion.

#### Bayesian regression on concentration parameters

To model the effects of bicarbonate and calcium on the subpopulation composition, we expressed the linear predictor *η*_*i,k*_ for each well *i* and subpopulation *k* as:

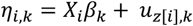

Where *X*_*i*_ denotes the row of the design matrix corresponding to assay well *i* (a 1 × 4 vector of predictors in this case), and *β*_*k*_ is a 4 × 1 column vector of fixed effects describing how each predictor shifts proportions of subpopulation (*k*). *u*_*z*[*i*],*k*_ is a random intercept for mouse *z* from which the [*i*] sample was collected, accounting for baseline differences among animals due to biological variation. The linear predictors for all subpopulations in assay well *i* were collected in the vector *η*_*i*_ = (*η*_*i* 1_, …, *η*_*i,K*_ ) . These were mapped to subpopulation probabilities *p*_*i*_ using a softmax link function:

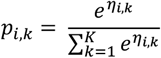

this ensures each *p*_*i,k*_ > 0 and ∑_*k*_ *p*_*i,k*_ = 1.

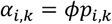

Where *ϕ* > 0 is a global concentration (intensity) parameter shared across assay wells and *p*_*i*_ =(*p*_*i*,1_, …, *p*_*i,K*_) is a probability vector on the K-simplex (i.e., a *K* ™ 1 dimensional space that represents all possible combinations of subpopulation proportions that sum to one). Under this parameterization, *p*_*i*_ represents the expected subpopulation proportions for assay well *i*, and *ϕ* controls the level of well-to-well variation.

#### Priors, standardization, and interpretation

To stabilize estimation and conservatively reflect weak prior information, we used the following prior distributions in the PyMC implementation:

Fixed effects—for each predictor *j* = 1, …, *J* and subpopulation *k* = 1, …, *K*, the fixed-effect coefficients were assigned independent priors:

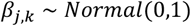

Random effects (mouse-level intercepts)—for each assay well *i* and subpopulation *k* = 1, …, *K*, a random intercept *u*_*z*[*i*],*k*_ was included to account for baseline differences among mice where *z*[*i*] indexes the mouse from which well *i* was obtained. The random effects were assumed to be independent across mice and subpopulations and were modeled as:

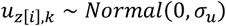

The standard deviation σ_*u*_, was assigned a *HalfNormal*(1) prior. Global concentration (overdispersion) hyperparameter:

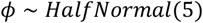

Which serves as a prior favoring moderate to large values while allowing the data to determine the extent of overdispersion. Continuous predictors (i.e., media bicarbonate, free calcium, and their interaction) were standardized to z-scores prior to fitting the model to place them on a comparable scale so that the priors on *β*_*k*_ corresponded to similar prior beliefs about effect sizes across predictors.

Because the mapping from the linear predictors to subpopulation proportions was nonlinear and coupled across subpopulations via the SoftMax link function, the coefficients *β*_*j,k*_ could not be interpreted directly as fold-changes in observed proportions. Instead, treatment effects are expressed in terms of posterior predicted proportions:

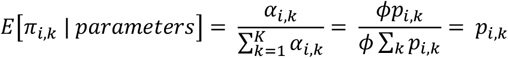

These expected values represent the model’s estimate of the ‘true’ underlying probability of each subpopulation for each treatment condition, accounting for both sampling noise and between animal variation.

For model fitting, the No-U-Turn Sampler (NUTS) from PyMC was used for Markov Chain Monte Carlo sampling^41^. Hamiltonian Monte Carlo sampling that NUTS employs was previously found to result in greater accuracy in compositional analysis of simulated datasets compared to other sampling methods^45^. Sampling was conducted with 4 independent Markov chains, each drawing 1000 posterior samples after 1000 tuning iterations. The target acceptance probability was 0.9. Convergence was assessed using the Gelman-Rubin statistic 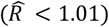 and effective sample size > 400 per parameter. Divergent transitions were not observed with these hyperparameter choices. Posterior summaries, raw proportion counts, medians, and 95% credible intervals were computed for all fixed and random effects.

## Results

### A 2D subculture assay for assessing signaling interaction effects during in vitro sperm capacitation

We first sought to develop a culture assay system to measure average cell physiological state changes in response to controlled signaling inputs, we employed 2D microtiter fluorometric assays (Figure 1A). The assays enabled assessment of changes in response (e.g., intracellular calcium, cell viability, etc) to 1) increasing concentrations of bicarbonate and calcium, 2) the change over time, and 3) the interaction effects between these two predictors. These measurements revealed that bicarbonate stimulation promotes calcium uptake in a time dependent manner using radiometric Indo-1 AM (K_d_ ∼230nM) (Fig1B). Increasing extracellular free calcium concentration increased Indo-1 ratio (i.e., bound/unbound) but did not exhibit kinetic effects, indicating that a relatively greater average baseline intracellular calcium concentration can be forced by increasing free calcium in the media, but that bicarbonate is still required to facilitate uptake. The kinetics and endpoint Indo-1 ratio were approximately matched between positive control conditions (calcium ionophore ionomycin) and a bicarbonate + calcium condition (10mM and 1.3 mM respectively) (Figure 1B). Cell viability was simultaneously monitored using live-cell impermeable (cationic) ethidium homodimer-1, which transitions from weakly fluorescent to strongly fluorescent when it binds DNA (Figure 1C). Positive control included detergent treated (permeabilized) cells. Interestingly, treatment groups that included bicarbonate stimulated the greatest loss of cell viability over time. Together these results predict that calcium uptake is kinetically dependent on bicarbonate, and that bicarbonate is positively correlated with loss of cell viability.

**Figure 1.**
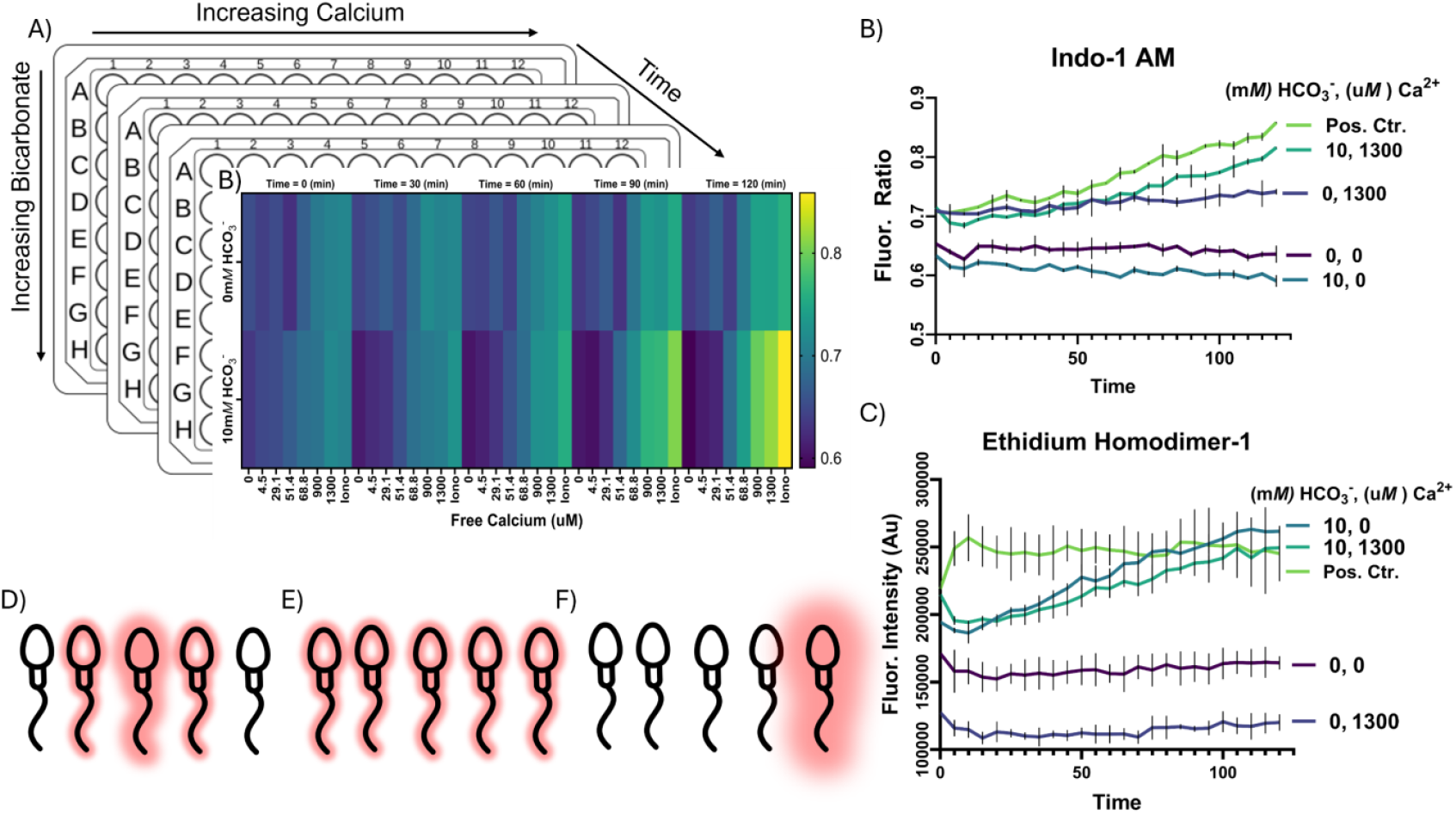
A 2D subculture assay for assessing signaling interaction effects during in vitro sperm capacitation. A) Diagram of a 96-well plate design allowing combinations of chemically ‘clamped’ free calcium and bicarbonate concentrations, each representing distinct microenvironments that sperm might encounter during the search for an egg. B) Kinetic plots showing Indo-1 ratios over time for sperm exposed to 0 and 10 mM bicarbonate concentrations over a corresponding range of chemically ‘clamped’ extracellular calcium, demonstrating the interaction effects apparent in bicarbonate signaling control of intracellular calcium uptake. C) Viability measurements using ethidium homodimer-1, a live-cell impermeable dye, showing how external conditions affect sperm viability over time. D–F) Conceptual examples of possible sperm subpopulation heterogeneity in responses to external signals wherein the same total signal can be reflected in completely different underlying distributions: D) A symmetric (Gaussian-like) distribution. E) A uniform response. F) A power distribution wherein 80% of the signal is localized among only 20% of the cells.

Notably, there is a critical limitation of interpreting bulk cell-population measurements common to this sort of approach—i.e., the inability to resolve how individual cells within a population respond to changes in the predictor. The fluorescent signal in both cases (Indo-1 AM and Ethidium homodimer-1) is the sum signal of the entire cell population. The shape of the distribution that signal is drawn from cannot be assumed because the fluorescence intensity may be distributed among the cells in any number of different ways while still retaining the same mean fluorescence intensity. As examples the distribution of signal could follow: 1) an exponential with a small number of sperm being very brightly fluorescent (Figure 1D), 2) a uniform distribution in which all of the cells exhibit similar intensities (Figure 1E), or 3) a power law distribution with ∼20% of the sperm accounting for ∼80% of the signal (Figure 1F). There are many other possible distributions not discussed here, and importantly, the shape of signal distribution could be different for various experimental treatment conditions. Thus, this type of ‘bulk’ measurement assay simply cannot distinguish between these very different scenarios and has a limited utility as a result - a fact which is true for all similar assays that do not make measurements with single-cell resolution such as western blotting, mass spectrometry label-tracing analysis, etc.

### Gating strategy for live-intact cell identification

Flow cytometry measures particles at or above the size of the nano length scale, including media components, cellular fragments, and intact cells—necessitating extensive controls for live-cell detection. To improve the accuracy of cell identification, all replicates of spectral flow data included cell-free and single-stain controls. When samples were treated with digitonin to permeabilize the cells resulting in loss of intracellular contents and a shift in refractive index, the density of scanned events shifted notably (Fig. 2A, B), consistent with membrane permeabilization and increased detection of subcellular material. In untreated samples gated for live cells, fewer total events were detected (Fig. 2C, D). After gating for single cells, all remaining events were positively stained for TO-PRO-3, indicating loss of membrane integrity and widespread cell death (Fig. 2E, F).

**Figure 2.**
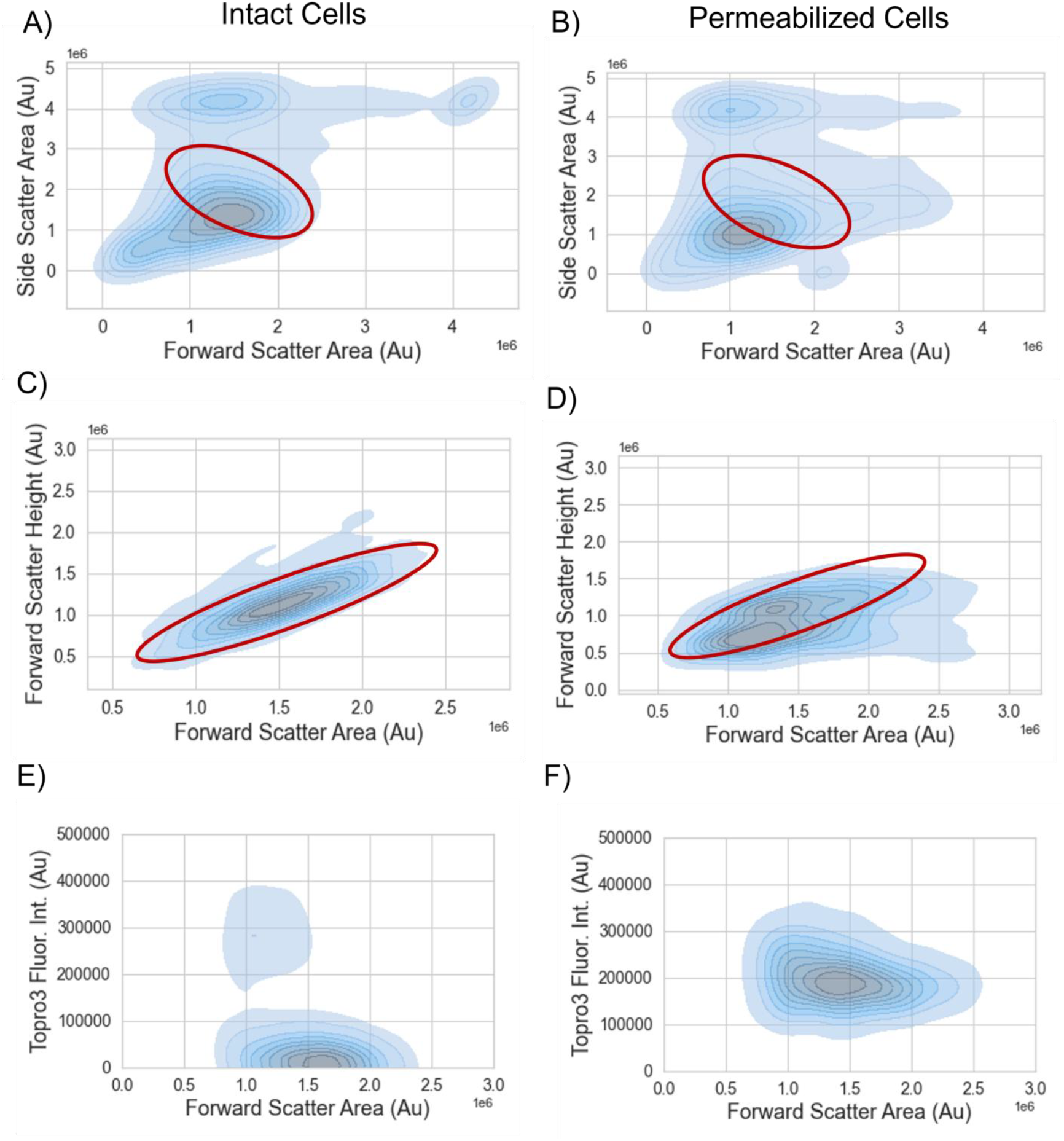
Gating strategy for live-intact cell identification. A) Representative forward and side scatter area contour plot showing all detected particles, including media, debris, and cells. B) Forward and side scatter area contour plot showing increased detection of small particles and cell debris after digitonin permeabilization. The intact cell gate remains highlighted in red. (C) Doublet exclusion using forward scatter area (FSC-A) versus height (FSC-H), used to aggregates from the analysis for intact cells. (D) Doublet exclusion using forward scatter area (FSC-A) versus height (FSC-H) to remove aggregates from permeabilized cell samples. (E) TO-PRO-3 viability staining revealed live (membrane-intact) cells as TO-PRO-3–negative, confirming a high proportion of intact sperm in this gate. (F) TO-PRO-3 staining labeling permeabilized whole cells with high TO-PRO-3 fluorescence signal, confirming membrane disruption and allowing clear discrimination between live-intact cells and dead-permeabilized cells.

Mouse sperm are delicate cells and will ‘decapitate’ easily when the haploid nucleus separates from the midpiece. To account for this common cell fragmentation pattern in the gating scheme, sperm were stained with Hoechst 33342 and subjected to repeated shear stress (mechanical vortex) for 5 minutes to disrupt their structure. Hoechst staining in combination with forward and side scatter measurements facilitated identification of gating regions that contained decapitated cells (not shown).

### Validation of multiplex dye localization

To examine capacitation responses under controlled conditions, the 2D fluorometric assay was modified to include three bicarbonate concentrations (0, 5, and 10 mM) and four chemically ‘clamped’ extracellular free calcium concentrations (0, 87.45, 660, and 1300 μM). To validate that the spectral flow cytometry measurements accurately reflect biological interpretation, sperm were stained with the full dye multiplex group: Indo-1 AM, PNA lectin–FITC, and TO-PRO-3 and imaged using laser scanning confocal microscopy. Indo-1 localizes to the head and midpiece, consistent with regions of high calcium signaling activity. PNA lectin-FITC weakly labeled the entire sperm surface but intensely labeled the acrosomal region in reacted sperm consistent with previous reports^36^ (Figure 3). Live-cell impermeable TO-PRO-3 selectively stains nuclear DNA. Together, the observed staining patterns confirm the expected spatial distribution of each dye and support their use for downstream classification in our flow cytometry assays.

**Figure 3.**
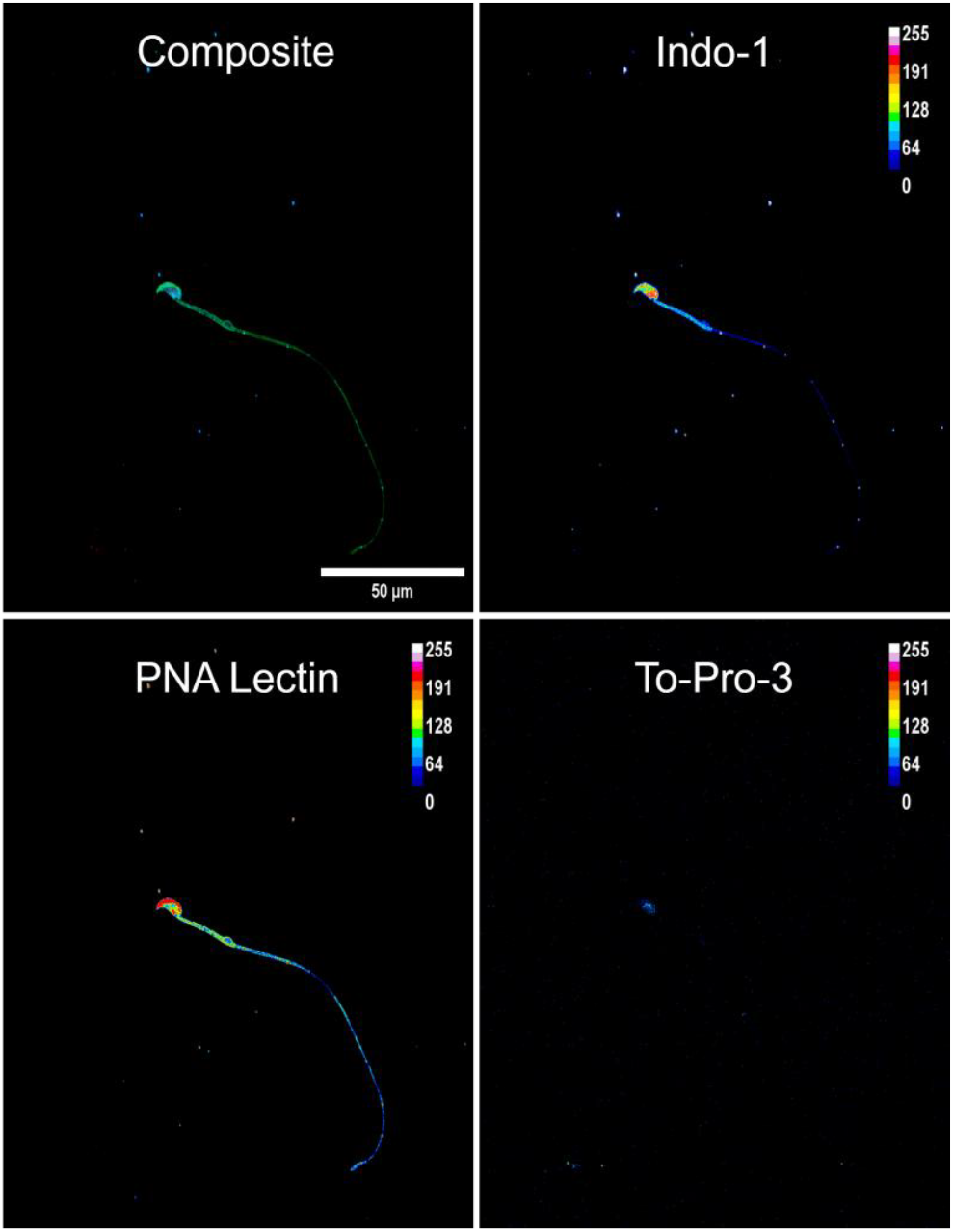
Validation of multiplex dye localization. Indo-1 localizes to the nucleus and midpiece, FITC conjugated peanut agglutinin (PNA) lectin labels whole cell with stronger relative signal at the acrosomal cap once exposed, and TO-PRO-3 fluorescence indicates nuclear DNA in non-viable (dye permeable) cells. Scale bar = 50 µm. Calibration bar = 16-color for 8-bit encoding.

### Classification of live sperm into subpopulations

Qualitative manual gating is common among flow cytometry studies. Here, we employed statistical classification methods to reduce potential bias and automate the process of classifying sperm subpopulations. Given the fluorescent dye compliment that we used, there were four possible subpopulation classes (Figure 4A): 1) Live/Acrosome Unreacted (L_UR_), 2) Live/Acrosome Reacted (L_R_), 3) Dead/Acrosome Unreacted (D_UR_), and 4) Dead/Acrosome Reacted (D_R_). To scale the single cell responses to ground truth effects, positive and negative control conditions were used in every biological replicate tested, with calcium ionophore (A23187) as an acrosomal exocytosis positive control and permeabilizing detergent (digitonin) as a cell death positive control. The positive controls contained exogenous clamped 1.3 mM CaCl_2_ and 10 mM HCO_3_. The negative controls contained no exogenous CaCl_2_ or HCO_3_. The ToPro3 and PNA lectin-FITC responses were smoothed using kernel density estimates to approximate a probability density function for each intensity (Figure 4B, C). Representative probability density plots with KDE’s demonstrated distinct probability densities for positive control condition. The probability density estimates were then used in conjunction with a naive Bayes classifier to assign every measured cell in the treatment groups to either positive or negative control classes. Joint class distributions were assigned to one of each of the four subpopulations outlined above.

**Figure 4.**
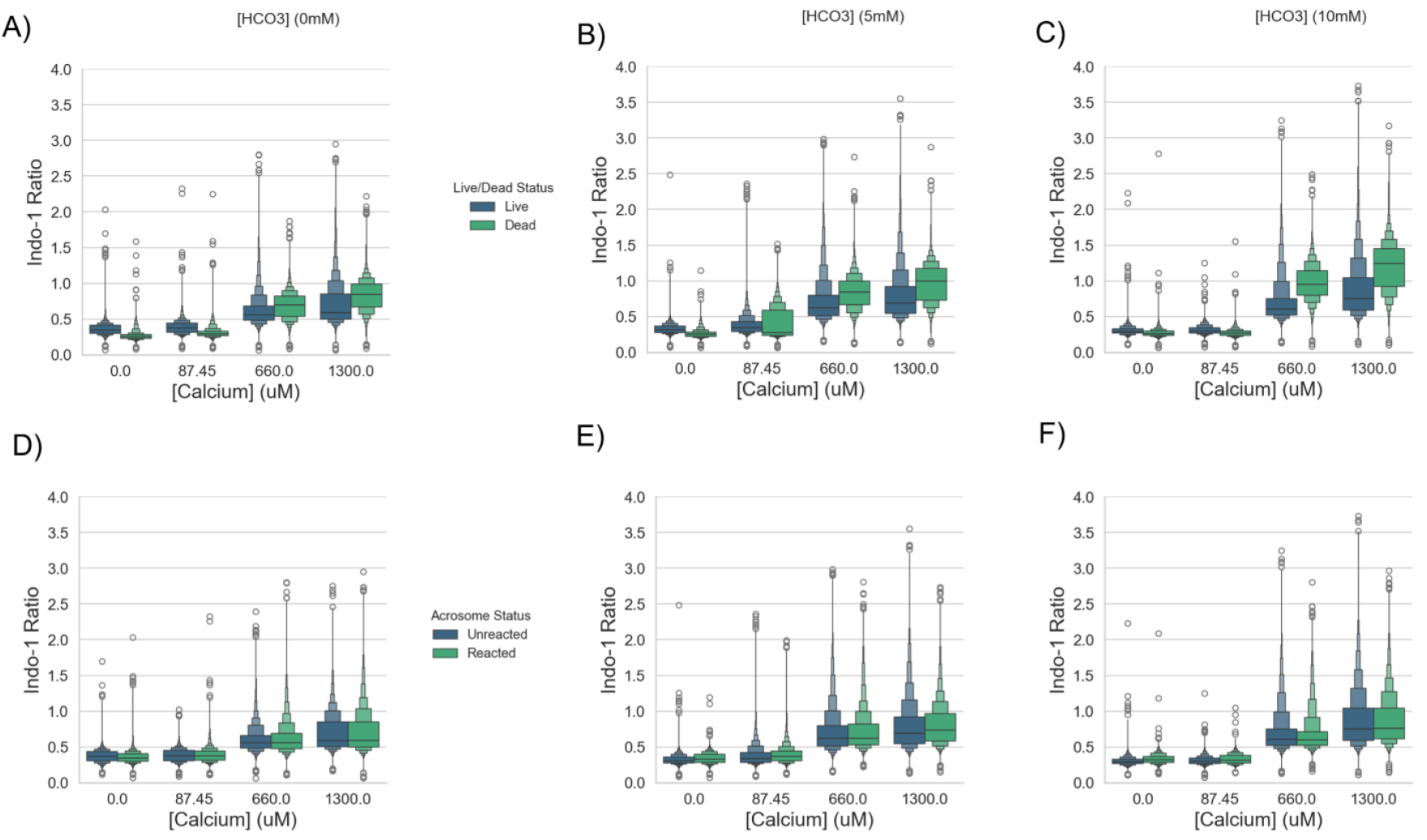
Effect of extracellular calcium and bicarbonate on cell viability and acrosome reaction. (A-C) Indo-1 ratio (intracellular calcium) as a function of chemically ‘clamped’ extracellular calcium (in µM) for classes grouped by live/dead status. Extracellular bicarbonate increasing from left to right (0-10 mM). Stacked Boxen plots show quantiles for visualization of underlying distribution shapes. Plots include data pooled from N=6 mice, totaling approximately 4.62 million cells. (D-F) Indo-1 ratio (intracellular calcium) as a function of chemically ‘clamped’ extracellular calcium (in µM) for classes grouped by acrosome reacted status. Only live cells were included in plots. Extracellular bicarbonate increasing from left to right (0-10 mM). Plots include live-cell data pooled from N=6 mice, totaling approximately 3.69 million cells. Outliers indicated by circles (1.5x interquartile range).

The observed off-target labeling of PNA lectin-FITC (described in the previous section) complicated the spectral interpretation due to signal that increases within a relatively small dynamic range during acrosomal exocytosis. However, the shift in probability density between positive and control conditions consistently revealed a high signal intensity region with greater relative probability density (Figure 4B). Additionally, KDE’s for ToPro3 gave very good separation between positive and negative control conditions, confirming membrane-compromised (non-viable) cells with distinct nuclear fluorescence could be detected (Figure 4C).

The extent to which change in intracellular calcium controls acrosomal exocytosis during capacitation is an ongoing topic of debate. We examined the distribution of Indo-1 ratio among in a representative population of cells from a single mouse to qualitatively assess the effect of microenvironmental conditions on intracellular calcium (Figure 4D). Indo-1 ratios of all subpopulations were relatively similar at low extracellular calcium concentrations. High calcium x bicarbonate treatment shifted the distribution of subpopulations toward a low-density subpopulation of high intracellular calcium sperm dominated largely by dead cells. The live, acrosome unreacted subpopulation remained the bulk of the total population under all treatment conditions. Together, these data demonstrate that cauda epididymal sperm from the same animal can respond to a range of signaling impulse intensities by shifting the composition of cell subpopulations. Under each treatment condition, the subpopulation distributions exhibited differing degrees of heterogeneity.

### Effect of extracellular calcium and bicarbonate on cell viability and acrosome reaction

To extend our previous qualitative analysis of within-mouse indo-1 ratio distributions for each subpopulation, we examined the class distributions of sperm phenotypes using pooled data from N=6 mice (∼4.62 million sperm) and plotted each cell’s Indo-1 ratio against the experimental treatment conditions (bicarbonate x calcium) for each mouse. Using a KDE-Bayes classifier, individual sperm were classified as either live or dead based on TO-PRO-3 staining patterns compared with positive and negative controls. Data did not appear to be symmetrically distributed, indicating that they would not fulfill assumption criteria for statistical designs that are typically implemented in flow cytometry studies. At low calcium concentrations, median Indo-1 ratios were greater in live cells than dead cells (Figure 5 A-C). Interestingly, this relationship flipped, and median Indo-1 ratios were greater in dead cells than live cells at high calcium concentrations (0.66 and 1.3 mM). For all conditions, Indo-1 ratios were positively correlated with increasing extracellular calcium and bicarbonate, but the effect was larger among dead cells likely because dead cells equilibrate calcium passively. Next, we examined the distributions of cells classed as acrosome reacted or unreacted in live cells pooled from N=6 mice (∼3.69 million live sperm). Surprisingly, no correlation was observed between Indo-1 ratio and acrosome reacted status in any of the treatment conditions (Figure 5D-F).

**Figure 5.**
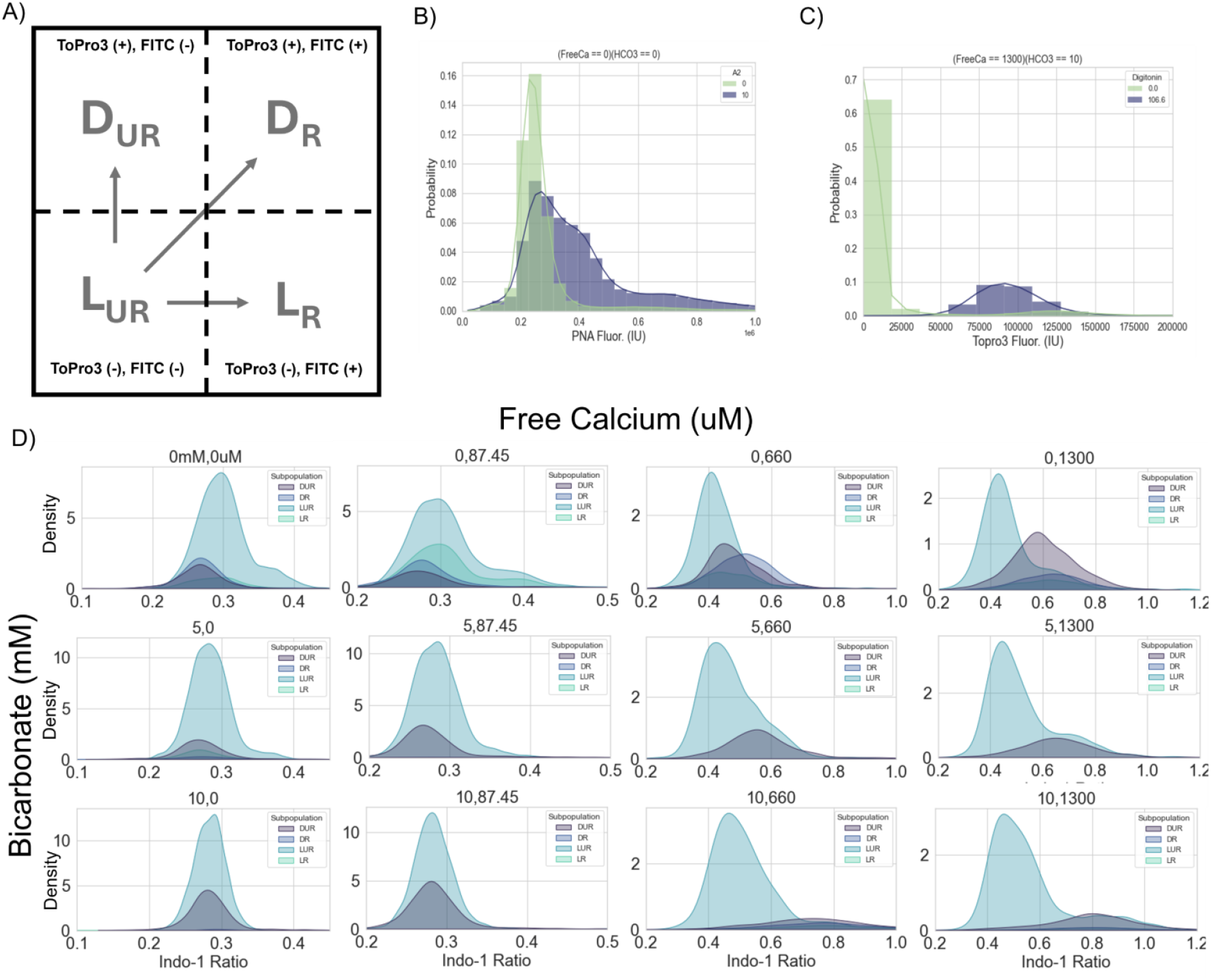
Classification into subpopulations. (A) diagram demonstrating possible cell subpopulation classifications for the response variables measured in this study and corresponding fluroscent dye interpretations. L_UR_= Live/Unreacted, L_R_= Live/Reacted, D_UR_= Dead/Unreacted, and D_R_= Dead/Reacted. Notably, each class represents an *irreversible* physical process. (B,C) Representative probability density plots for positive and negative controls conditions demonstrating distinct probability distributions (fitted to kernel density estimates) for each condition class (i.e., live/dead, acrosome reacted/unreacted). (B) PNA-FITC fluorescence for positive control (10 µM A23187) and negative control (ionophore free, low calcium, low bicarbonate) conditions. (C) TO-PRO-3 fluorescence for positive control (106.6 µM digitonin) and negative control (detergent free, low calcium, low bicarbonate) conditions. (D) Representative kernel density estimate (KDE) plots for each classified subpopulation Vs. corresponding intracellular calcium (Indo-1 ratio). Plots organized by each pairwise combination of extracellular bicarbonate (mM) and calcium (µM).

### Dirichlet-Multinomial hierarchical regression model of subpopulation composition as an adaptive response to microenvironmental conditions

Rather than treating subpopulations as statistically independent, we analyzed the conditional changes in subpopulation distributions using a hierarchical Bayesian regression model. To examine the subpopulation distributions among each treatment condition, we plotted the observed sampling proportions (Figure 6A-C). In mouse sperm, bicarbonate activates soluble adenylate cyclase, which increases calcium permeability through cAMP dependent interactions with the sperm specific cation channel family (CatSper)^8^. Conditional effects on the proportions of Live acrosome unreacted and all dead sperm were relatively stable across conditions. Interestingly, the highest proportion of live acrosome reacted cells was observed at low extracellular calcium concentrations independent of extracellular bicarbonate concentration (Figure 6A-C).

**Figure 6.**
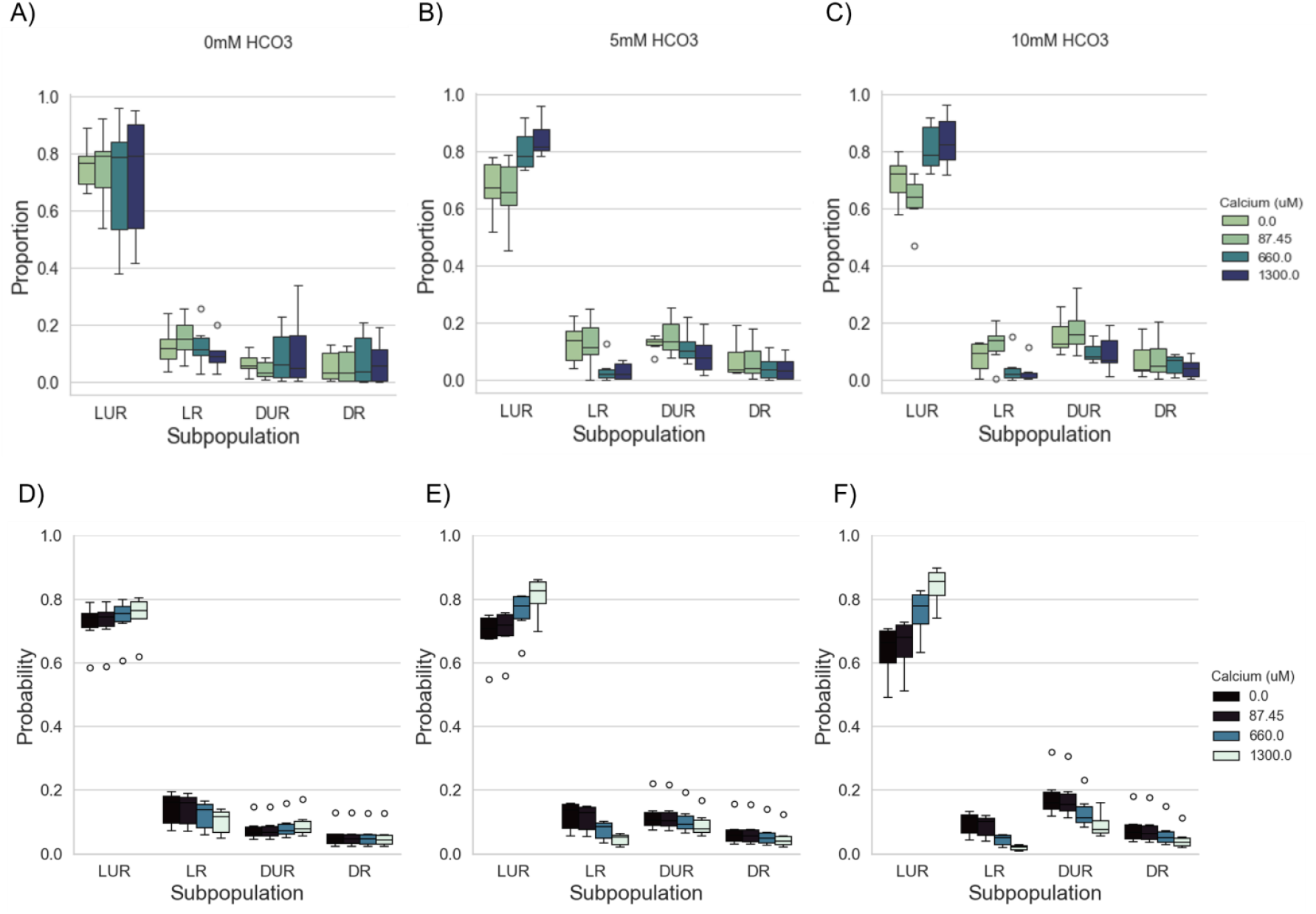
Dirichlet-Multinomial hierarchical regression model of subpopulation composition as an adaptive response to microenvironmental conditions. (A-C) Measured proportions of subpopulation classes across each of the treatment conditions (*HCO*_*3*_ *x Calcium*) for N=6 mice. Outliers indicated with filled diamonds (1.5x IQR). (D-F) Mean posterior probability estimates of the underlying multinomial sampling probabilities of subpopulation compositions for each treatment condition. Outliers indicated by circles (1.5x IQR). Bicarbonate concentrations for each treatment indicated at the top of the panel. Corresponding model credible intervals are provided in table 7.

The posterior predictions from our hierarchical regression model indicate a few interesting patterns regarding expected subpopulation probabilities under various microenvironmental signaling conditions (Table 1). First, the expected proportion of live unreacted (L_UR_) sperm was inversely related to both bicarbonate and calcium concentration reflected in an ∼7-20% difference in L_UR_ proportion between calcium/bicarbonate free conditions and those of high concentrations. Second, the subpopulation of dead reacted (D_R_) cells did not appreciably change in a patterned way among any of the conditions, indicating that acrosomal exocytosis likely does not predispose sperm to an increased probability of death. Finally, the predicted proportion of dead unreacted (D_UR_) sperm indicated that the ratio of bicarbonate to free calcium may be important – with apparent toxicity when bicarbonate was present at low relative calcium concentrations particularly at higher bicarbonate concentration (i.e. 10mM).

**Table 1.** Summary of median posterior probabilities *E*(π_*i,k*_|*parameters*). Expected multinomial sampling probabilities for each subpopulation under each treatment condition. Accompanying 95% credible intervals (*q*2.5%, *q*97.5%). N= 5 mice.

| [HCO<sub>3</sub>]<br>(mM) | [Ca<sup>2+</sup>]<br>(μM) | Live Unreacted<br>(L<sub>UR</sub>) | Live Reacted (L<sub>R</sub>) | Dead Unreacted<br>(D<sub>UR</sub>) | Dead Reacted<br>(D<sub>R</sub>) |
| --- | --- | --- | --- | --- | --- |
| 0 | 0 | 0.71 (0.58, 0.81) | 0.14 (0.08, 0.24) | 0.07 (0.04, 0.13) | 0.05 (0.02, 0.08) |
| 0 | 87.45 | 0.72 (0.59, 0.81) | 0.14 (0.08, 0.24) | 0.07 (0.04, 0.13) | 0.05 (0.03, 0.10) |
| 0 | 660 | 0.73 (0.61, 0.82) | 0.12 (0.07, 0.20) | 0.08 (0.05, 0.15) | 0.05 (0.03, 0.09) |
| 0 | 1300 | 0.74 (0.62, 0.84) | 0.10 (0.05, 0.18) | 0.09 (0.05, 0.17) | 0.05 (0.02, 0.10) |
| 5 | 0 | 0.68 (0.56, 0.78) | 0.12 (0.07, 0.19) | 0.12 (0.07, 0.19) | 0.06 (0.04, 0.11) |
| 5 | 87.45 | 0.69 (0.57, 0.79) | 0.11 (0.06, 0.18) | 0.12 (0.07, 0.20) | 0.06 (0.04, 0.11) |
| 5 | 660 | 0.75 (0.64, 0.83) | 0.07 (0.04, 0.12) | 0.10 (0.06, 0.18) | 0.05 (0.03, 0.10) |
| 5 | 1300 | 0.80 (0.70, 0.87) | 0.05 (0.02, 0.08) | 0.09 (0.05, 0.17) | 0.04 (0.02, 0.08) |
| 10 | 0 | 0.63 (0.49, 0.74) | 0.10 (0.05, 0.16) | 0.18 (0.10, 0.29) | 0.07 (0.04, 0.14) |
| 10 | 87.45 | 0.64 (0.51, 0.76) | 0.09 (0.04, 0.14) | 0.17 (0.10, 0.28) | 0.07 (0.04, 0.13) |
| 10 | 660 | 0.75 (0.63, 0.84) | 0.04 (0.02, 0.07) | 0.13 (0.07, 0.23) | 0.06 (0.03, 0.10) |
| 10 | 1300 | 0.84 (0.73, 0.90) | 0.02 (0.007, 0.04) | 0.09 (0.05, 0.17) | 0.04 (0.02, 0.09) |

## Discussion

This report reveals that mouse sperm capacitation is characterized by substantial and structured phenotypic heterogeneity, even among cells originating from the same male and exposed to identical microenvironmental conditions. Using spectral flow cytometry to assess sperm phenotype distributions with single-cell resolution, combined with a controlled two-dimensional titration of bicarbonate and extracellular free calcium, we demonstrate that sperm do not respond uniformly to capacitating signals. Instead, they redistribute across distinct physiological subpopulations in a manner that depends nonlinearly on the impulse strengths of the signaling inputs.

The findings are consistent with earlier work showing variability in calcium permeability^16,17^, membrane potential^18,33^, tyrosine phosphorylation^20^, and acrosome reaction propensity^33,46–48^. The diversity of responses observed, and their sensitivity to microenvironmental conditions emphasizes that capacitation is not a simple synchronized transition, but a population-level stochastic process shaped by complex biochemical determinants that vary among large numbers of cells. For these reasons, ensemble averaging in sperm or semen samples under the assumption that sperm vary independently is unlikely to carry strong predictive value. This is important because the dynamic structure of sperm subpopulations is likely to be biologically consequential for fertility and may provide useful information for the quantitative prediction of fertilizing potential from semen analysis.

The interaction between bicarbonate and extracellular calcium revealed several unexpected and informative patterns. Although both ions are required for canonical capacitation signaling^3–6,8^, the highest proportions of live, acrosome-reacted sperm were observed under relatively low extracellular calcium and low bicarbonate. Though somewhat surprising, this finding is consistent with a previous report of biphasic effects on tyrosine phosphorylation induced by controlled extracellular calcium in the presence of a chelating agent^49^. Increasing either stimulus elevated intracellular calcium but simultaneously increased cell death, producing subpopulations dominated by non-viable, calcium-loaded cells. These findings align with reports that intracellular calcium elevation is necessary but not sufficient for spontaneous acrosomal exocytosis^5,8,9^, and that excessive calcium influx can compromise viability. Although notably, this effect may also result from non-viable cells inability to actively maintain a low intracellular calcium potential. The apparent associations between bicarbonate, calcium load, and acrosome reaction status in live cells suggests that capacitating stimuli may push many sperm toward non-functional endpoints, reducing the fraction capable of fertilization. Interestingly, bicarbonate appeared to be conditionally toxic to mouse sperm when extracellular calcium was low.

The comparison between bulk cell spectrofluorometric measurements and single-cell flow cytometry further highlights the importance of measuring sperm phenotypes with single-cell resolution. Population-averaged fluorescence masked fundamentally different underlying distributions, underscoring longstanding concerns about interpreting capacitation solely from ensemble measurements^1,15^. Single-cell approaches are particularly important because the fertilizing potential of an ejaculate is best understood as a statistical property of many cells, not as the behavior of a hypothetical “best” sperm implied by ensemble averaging.

The Dirichlet–multinomial hierarchical regression model provides a principled framework for quantifying how treatments reshape the entire subpopulation composition while accounting for multinomial sampling noise and biological variation among biological replicates. By modeling conditional changes in the expected proportions of live unreacted, live reacted, dead unreacted, and dead reacted cells - this approach avoids the interpretive pitfalls of treating subpopulations independently. The finding that between-male variation in baseline subpopulation structure is substantial is also consistent with previous reports of high within- and between-individual variability in sperm functional phenotypes among human males^22,23^. The resulting predictive probability surfaces provide a quantitative representation of sperm subpopulation responses to extracellular cues and complement mechanistic models that attribute capacitation heterogeneity to variable ion channel expression or signaling kinetics^17,19,21^.

The results also relate to broader evolutionary questions about sperm number and diversity. Many species produce sperm in numbers far exceeding those required for fertilization, but fertility also collapses at ‘low’ sperm counts that may still consist of many thousands or even millions of cells^24^. Heterospermic insemination experiments have shown that phenotypic heterogeneity contributes to adaptive advantages in uncertain reproductive environments such as female reproductive tracts of hybridizing species^27,50^. Our findings support the hypothesis that sperm populations hedge against uncertainty by distributing cells across multiple physiological states that are sensitive to mechanistic signals through the statistical variability of sperm subpopulations. In vivo, only a small minority of sperm that reach the oviduct display an intact acrosome^20,46^, and the timing of capacitation appears to align with anticipation of egg availability^11^. A heterogeneous distribution of phenotypes may therefore maximize the likelihood that some sperm encounter the egg in an appropriately primed state.

Several limitations should be noted for this study. First, interpretation of acrosomal status is constrained by the modest dynamic range of PNA-FITC in live cells and its susceptibility to off-target binding^36^. Though previous reports have shown that live cell measurement of acrosome reaction using PNA-FITC correlates well with measurements of acrosomal exocytosis in acrosin-GFP transgenic mice^47,48^, the proportion of AR positive cells following calcium ionophore stimulation tends to be comparatively lower with use of PNA-FITC, indicating that the AR subpopulation is not completely captured with this method. Despite that limitation, our results highlight that distinguishing live from dead cells is essential, and that methods employing organic extraction with PNA-FITC significantly overestimate fertility competent cells because they cannot make the necessary distinction between these subpopulations^51^. Second, although bicarbonate and calcium titrations capture key components of capacitation signaling, in vitro conditions cannot fully reproduce the biochemical and physical complexity of the oviductal environment, particularly fluid rheology, chemokinesis, and structural constraints^12–14^. Third, the design in this report is focused on a single timepoint rather than a full temporal trajectory; measuring time dynamics of subpopulation transitions between capacitation states will require separate time-lapse flow cytometry experiments.

## Conclusion

In summary, this work demonstrates that sperm capacitation can be modeled as a stochastic, cell population-level adaptive process shaped by structured physiological heterogeneity. Importantly, the adaptability of the whole cannot exceed the adaptability of the component subpopulations. Understanding how subpopulation compositions vary under physiologically relevant microenvironmental signaling conditions will enable modeling of fertilizing potential as a function of the whole ejaculate, rather than relying on assumptions about idealized individual sperm phenotypes (i.e., the ‘best’ sperm). By integrating large-scale single-cell flow cytometry with a hierarchical statistical framework for modeling sperm subpopulation compositions, we provide a quantitative foundation for analyzing and predicting how changing microenvironmental conditions redistribute cells among functional subpopulations. Future work should incorporate time-resolved measurements, physiologically relevant media conditions, and functional assays linking subpopulation structure to measurable fertility outcomes.

## Author Contributions

BB, AH, AC, AW, MA, DH, DB, PV, and CAS designed experiments and collected data. BB and CAS authored and edited the report. All co-authors reviewed and approved the work.

## Acknowledgements

None

## References

1. Gervasi, M. G. & Visconti, P. E. Chang’s meaning of capacitation: A molecular perspective. Molecular Reproduction and Development vol. 83 860–874 Preprint at 10.1002/mrd.22663 (2016).

2. Schmidt, C. A. et al. Pyruvate modulation of redox potential controls mouse sperm motility. Dev Cell https://doi.org/10.1016/j.devcel.2023.11.011 (2023) doi:10.1016/j.devcel.2023.11.011.

3. Santi, C. M. et al. The SLO3 sperm-specific potassium channel plays a vital role in male fertility. FEBS Lett 584, 1041–1046 (2010).

4. Nishigaki, T. et al. Intracellular pH in sperm physiology. Biochemical and Biophysical Research Communications vol. 450 1149–1158 Preprint at 10.1016/j.bbrc.2014.05.100 (2014).

5. Hess, K. C. et al. The ‘soluble’ adenylyl cyclase in sperm mediates multiple signaling events required for fertilization. Dev Cell 9, 249–259 (2005).

6. Visconti, P. E. et al. Capacitation of mouse spermatozoa I. Correlation between the capacitation state and protein tyrosine phosphorylation. Development 1129–1137 (1995).

7. Visconti, P. E. et al. Capacitation of mouse spermatozoa II. Protein tyrosine phosphorylation and capacitation are regulated by a cAMP-dependent pathway. Development 1139–1150 (1995).

8. Tateno, H. et al. Ca2+ ionophore A23187 can make mouse spermatozoa capable of fertilizing in vitro without activation of cAMP-dependent phosphorylation pathways. Proc Natl Acad Sci U S A 110, 18543–18548 (2013).

9. Suarez, S. S. Control of hyperactivation in sperm. Human Reproduction Update vol. 14 647–657 Preprint at 10.1093/humupd/dmn029 (2008).

10. Hirohashi, N. & Yanagimachi, R. Sperm acrosome reaction: Its site and role in fertilization. Biology of Reproduction vol. 99 127–133 Preprint at 10.1093/biolre/ioy045 (2018).

11. Giojalas, L. C., Rovasio, R. A., Fabro, G., Gakamsky, A. & Eisenbach, M. Timing of sperm capacitation appears to be programmed according to egg availability in the female genital tract. Fertil Steril 82, (2004).

12. Zaferani, M. & Abbaspourrad, A. Biphasic Chemokinesis of Mammalian Sperm. Phys Rev Lett 130, (2023).

13. Tung, C. K. et al. Fluid viscoelasticity promotes collective swimming of sperm. Sci Rep 7, (2017).

14. Schoeller, S. F., Holt, W. V. & Keaveny, E. E. Collective dynamics of sperm cells: Collective dynamics of sperm cells. Philosophical Transactions of the Royal Society B: Biological Sciences vol. 375 Preprint at 10.1098/rstb.2019.0384 (2020).

15. Martínez-Pastor, F. What is the importance of sperm subpopulations? Anim Reprod Sci 246, (2022).

16. Buffone, M. G., Doncel, G. F., Briggiler, C. I. M., Vazquez-Levin, M. H. & Calamera, J. C. Human sperm subpopulations: Relationship between functional quality and protein tyrosine phosphorylation. Human Reproduction 19, 139–146 (2004).

17. Luque, G. M. et al. Only a subpopulation of mouse sperm displays a rapid increase in intracellular calcium during capacitation. J Cell Physiol 233, 9685–9700 (2018).

18. Molina, L. C. P. et al. Membrane Potential Determined by Flow Cytometry Predicts Fertilizing Ability of Human Sperm. Front Cell Dev Biol 7, (2020).

19. Aguado-García, A., Priego-Espinosa, D. A., Aldana, A., Darszon, A. & Martínez-Mekler, G. Mathematical model reveals that heterogeneity in the number of ion transporters regulates the fraction of mouse sperm capacitation. PLoS One 16, (2021).

20. Ded, L., Hwang, J. Y., Miki, K., Shi, H. F. & Chung, J. J. 3D in situ imaging of female reproductive tract reveals molecular signatures of fertilizing spermatozoa in mice. Elife 9, 1–51 (2020).

21. Aldana, A., Carneiro, J., Martínez-Mekler, G. & Darszon, A. Discrete Dynamic Model of the Mammalian Sperm Acrosome Reaction: The Influence of Acrosomal pH and Physiological Heterogeneity. Front Physiol 12, (2021).

22. Steeman, T. J., Baro Graf, C., Novero, A. G., Buffone, M. G. & Krapf, D. High Variability in Human Sperm Membrane Potential over Time Can Limit Its Reliability as a Predictor in ART Outcomes. Biology (Basel) 14, (2025).

23. Keel, B. A. Within- and between-subject variation in semen parameters in infertile men and normal semen donors. Fertil Steril 85, 128–134 (2006).

24. Cohen, J. Why so many sperms? An essay on the arithmetic of reproduction. Sci Prog 57, 23–41 (1970).

25. Bell, A. D. et al. Insights into variation in meiosis from 31,228 human sperm genomes. Nature 583, 259–264 (2020).

26. Ollero, M. et al. Characterization of Subsets of Human Spermatozoa at Different Stages of Maturation: Implications in the Diagnosis and Treatment of Male Infertility. Human Reproduction vol. 16 (2001).

27. Pizzari, T., Worley, K., Burke, T. & Froman, D. P. Sperm competition dynamics: Ejaculate fertilising efficiency changes differentially with time. BMC Evol Biol 8, (2008).

28. Gur, Y. & Breitbart, H. Mammalian sperm translate nuclear-encoded proteins by mitochondrial-type ribosomes. Genes Dev 20, 411–416 (2006).

29. Brisard, B. M. et al. Modeling diffusive search by non-adaptive sperm: Empirical and computational insights. PLoS Comput Biol 21, (2025).

30. Schneider, C. A., Rasband, W. S. & Eliceiri, K. W. NIH Image to ImageJ: 25 years of image analysis. Nat Methods 9, 671–675 (2012).

31. Balestrini, P. A. et al. Seeing is believing: Current methods to observe sperm acrosomal exocytosis in real time. Molecular Reproduction and Development vol. 87 Preprint at 10.1002/mrd.23431 (2020).

32. Mckinney, W. Data Structures for Statistical Computing in Python. SciPy. 445, (2010).

33. Pedregosa, F. et al. Scikit-Learn: Machine Learning in Python. Journal of Machine Learning Research vol. 12 http://scikit-learn.sourceforge.net. (2011).

34. Virtanen, P. et al. SciPy 1.0: fundamental algorithms for scientific computing in Python. Nat Methods 17, 261–272 (2020).

35. Scott, D. W. Scott’s rule. Wiley Interdiscip Rev Comput Stat 2, 497–502 (2010).

36. Abril-Pla, O. et al. PyMC: a modern, and comprehensive probabilistic programming framework in Python. PeerJ Comput Sci 9, (2023).

37. Kumar, R., Carroll, C., Hartikainen, A. & Martin, O. ArviZ a unified library for exploratory analysis of Bayesian models in Python. J Open Source Softw 4, 1143 (2019).

38. Chen, J. & Li, H. Variable selection for sparse Dirichlet-multinomial regression with an application to microbiome data analysis. Annals of Applied Statistics 7, 418–442 (2013).

39. la Rosa, P. S. et al. Hypothesis Testing and Power Calculations for Taxonomic-Based Human Microbiome Data. PLoS One 7, (2012).

40. Harrison, J. G., Calder, W. J., Shastry, V. & Buerkle, C. A. Dirichlet-multinomial modelling outperforms alternatives for analysis of microbiome and other ecological count data. Mol Ecol Resour 20, 481–497 (2020).

